# Temporal variation in introgressed segments’ length statistics sheds light on past admixture pulses

**DOI:** 10.1101/2023.05.03.539203

**Authors:** Lionel N. Di Santo, Claudio S. Quilodran, Mathias Currat

## Abstract

Hybridization is recognized as an important evolutionary force, but identifying and timing admixture events between divergent lineages remains a major aim of evolutionary biology. While this has traditionally been done using inferential tools on contemporary genomes, the latest advances in paleogenomics have provided a growing wealth of temporally distributed genomic data. Here, we used individual-based simulations to generate chromosome-level genomics data for a two-population system and described temporal neutral introgression patterns under a single- and two-pulse admixture model. We computed three summary statistics aiming to inform the timing and number of admixture pulses between interbreeding entities: lengths of introgressed sequences and their variance within-genomes, as well as genome-wide introgression proportions. The first two statistics can confidently be used to infer inter-lineage hybridization history, peaking at the beginning and shortly after an admixture pulse. Temporal variation in introgression proportions provided more limited insights. We then computed these statistics on *Homo sapiens* paleogenomes and successfully inferred the hybridization pulse with Neanderthal that occurred approximately 40 to 60 kya. The scarce number of genomes dating from this period prevented more precise inferences, but the accumulation of paleogenomic data opens promising perspectives as our approach only requires a limited number of genomes.

## Introduction

Hybridization is defined as the inter-breeding between divergent lineages and is widespread in nature^1–3^. Where crossing between populations or species produces viable and fertile hybrids capable of backcrossing with parental individuals, genetic material can be transferred between lineages resulting in introgression. Previously underappreciated, recognition of the commonness of introgressive hybridization has stimulated research for both its detection and characterization. This includes interest in studying the direction, the frequency, the magnitude, the timing, the duration, and the mode of introgression (i.e., admixture pulses or continuous exchange of genetic material between lineages). While pulses of admixture are intuitively understood as discrete periods of interbreeding between divergent entities, it remains ambiguously characterized and poorly understood. Note that we will use the word “lineages” to refer to evolutionary units at different taxonomic levels, whether genus, species, subspecies or groups of individuals within species.

Over the last decades, multiple approaches have been designed to study introgresion^3–9^. Among phylogenetic-based techniques, the D-statistic, or ABBA-BABA test^10,11^, has been the most widely used to infer gene flow between lineages^12–15^. With the advent of next-generation sequencing technologies, large genomic datasets could be generated across many individuals, fostering the inclusion of coalescence modeling into the hybridization analytical toolbox. Combined with approximate Bayesian computation^16,17^, these methods have provided a powerful means to detecting among-lineage gene flow^18–20^. Importantly, these methods leverage present-time genetic information to infer historical gene flow, whereas patterns in introgression statistics evaluated temporally throughout ancient and modern samples may be able to provide a clearer picture of a lineage admixture history.

Following hybridization, genomic fragments from a donor lineage will enter a recipient lineage and only shorten with every backcrossing event due to recombination. Researchers have noticed the potential of this dynamic across generations to estimate the timing of admixture^6,8,21,22^ and, more recently, the duration of admixture^6^, as well as the strength of selection^4^. However, recombination may also challenge the inference of the number of admixture events, as time would most likely erase signatures of hybridization pulses within the genome (but see^9^). Analyzing the size distribution of introgressed fragments in time series genomic samples may be a promising avenue to accurately estimating the number and timing of multiple admixture events between lineages.

Genomic summary statistics including average length of introgressed fragments and within-genome variation in introgressed fragments’ lengths could be useful to estimate the number and timing of admixture pulses between hybridizing lineages. On the one hand, they can readily be estimated on single genomes. This is particularly important as the availability of ancient genomes may often be limited. On the other hand, both statistics are expected to vary substantially following gene flow between lineages. Indeed, we predict each pulse of admixture to increase the genome-wide average of the size of introgressed fragments before the action of recombination with passing generations ultimately erodes their size. Similarly, we predict the influx of longer fragments into a genomic background of shorter segments to increase within-genome variation in introgressed sequences’ lengths before the action of recombination homogenizes the size of introgressed fragments over time. Therefore, both average length of introgressed segments and within-genome variation in introgressed sequences’ lengths are expected to leave a perceptible imprint we can leverage, but they have yet to be described within a temporal context.

Recent advances in paleogenomics and the ensuing, ever-growing accessibility to ancient human genomes^23–26^, together with the considerable attention given to the history of archaic introgression into the genome of *Homo sapiens* during the past decade^27,28^, identifies our species as a candidate system to test this analytical framework. All non-African present-day humans share approximately 2% of their genomes with Neanderthal (*Homo neanderthalensis*)^10,29,30^. This result was first interpreted as evidence for a single pulse of admixture between *Homo sapiens* and Neanderthals in the Middle-East following the Out-Of-Africa (OOA) expansion^10^. However, with contrast to these findings, it was later found that East Asian populations may be more introgressed (by approximately 8-20%) than European populations^31–34^. This led scientists to hypothesize that *Homo sapiens* may have admixed with distinct Neanderthal populations through multiple pulses of hybridization during their shared evolutionary history^19,35,36^. Moreover, a continuous model of hybridization through space and time is also compatible with current Neanderthal introgression rate in humans^37,38^, although it is difficult to relate this model to a number of pulses. Ultimately, a decade exploring the overlapping evolutionary history of modern and archaic hominins has shed light on the complexity of studying inter-species hybridization, particularly when one wishes to evaluate the timing, number, and origin of hybridization pulses.

With this study, we first used genomic simulations to explore temporal patterns of introgressed segments’ length and its variation within individuals, in addition to introgression proportions, asking the question: can introgression summary statistics through time be leveraged to infer the number and timing of admixture pulses? Subsequently, we applied the knowledge gained from simulations to real empirical data, using the hybridization history between Neanderthals and *Homo sapiens* as a case study, to determine whether hypothesized pulses of admixture between these hominins could be identified based on proposed genomic summary statistics. Our study expands upon previous and contemporary work exploring and characterizing introgressive hybridization, focusing on temporal patterns of admixture tract lengths as well as a previously underexplored introgression statistics, within-genome variation in introgressed segments’ lengths, to explicitly infer the number and timing of hybridization pulses between divergent lineages.

## Results

### Admixture simulations

Temporal patterns of genomic summary statistics were generated by simulating the neutral genomic evolution through time of two populations connected by gene flow (a source - provider of migrants - and a sink - receiver of migrants - population) using individual-based modelling. Each population was populated by multiple individuals, whose genomic background were modeled as a single pair of homologous chromosomes experiencing recombination. Unadmixed individuals carry alleles typical of either the sink population or the source population. We considered two distinct models of admixture (Fig. 1), one simulating a single pulse of hybridization (referred to as the SP model, Fig. 1a) and one simulating two pulses of hybridization (referred to as the MP – multiple pulses – model, Fig. 1b). We also evaluated the effect of hybridization intensity on between-population exchange of genetic material by simulating both SP and MP with admixture rates of 0.03, 0.1, and 0.3 (hereafter referred to as scenarios). Following simulations, introgression within sink population was summarized using three population-scale genomic summary statistics: i) the average proportion of introgression (PI), ii) the average length of introgressed sequences (LIS), and iii) the average intra-individual variation in introgressed sequences’ lengths (VIS).

**Fig. 1.**
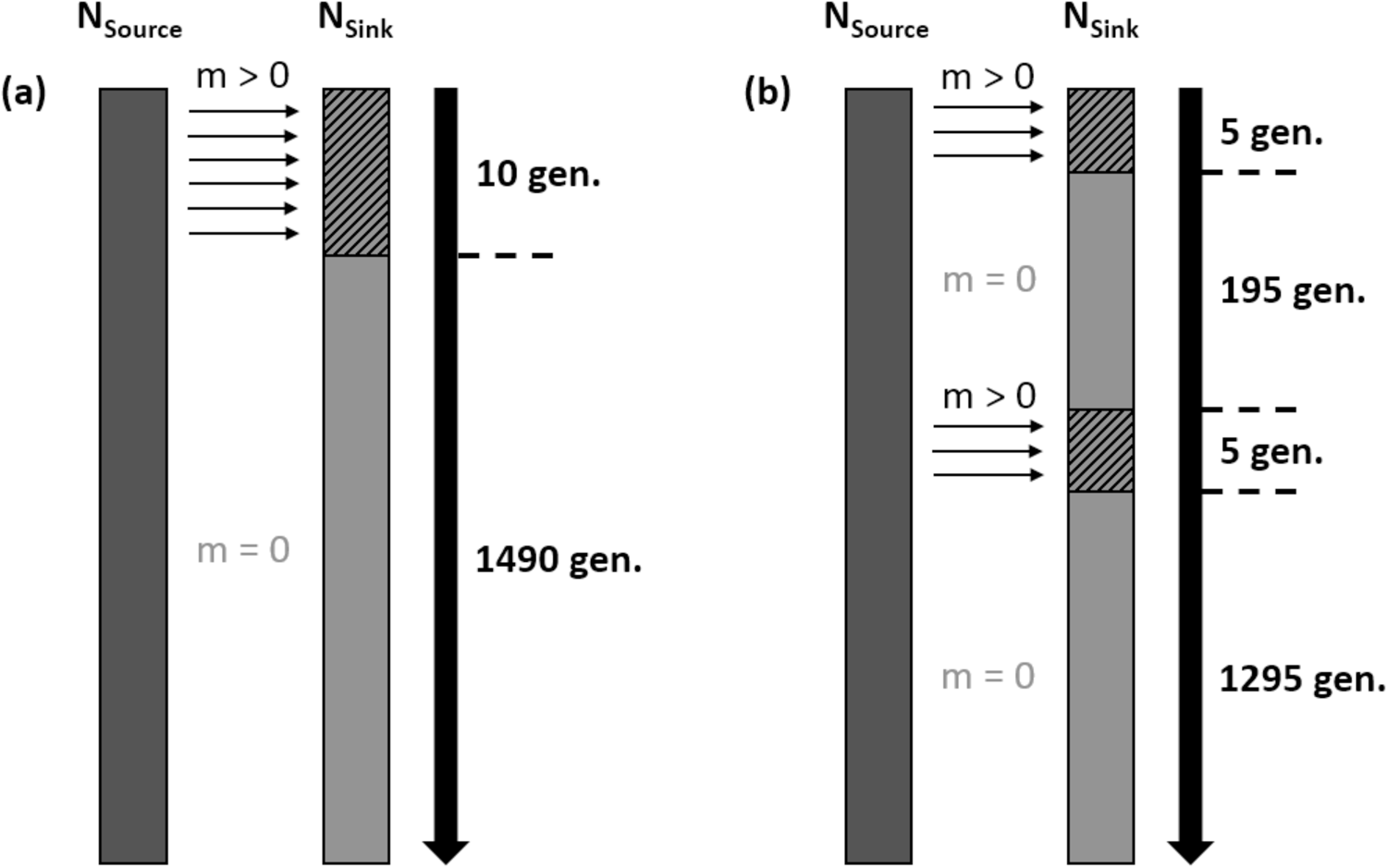
Schematics of admixture models, each simulated for a total of 1500 generations. (a) Single Pulse (SP) model. (b) Multiple Pulses (MP) model. Rectangles represent source (dark gray) and sink (gray) populations, with solid rectangle depicting generations in isolation and hashed rectangles depicting generations of admixture, small unidirectional arrows symbolize gene flow between populations and indicate its direction, and solid large arrows represent time with the number of generations spent in isolation or admixing given on the side. N_Source_/N_Sink_: Source/Sink initial population size (100 individuals), m: migration/admixture rate (either 0.03, 0.1, or 0.3 depending on the scenario simulated).

#### Proportion of introgression (PI)

Throughout generations, regardless of the admixture model and scenario of admixture rate simulated, average PI primarily increased while hybridization between source and sink populations was permitted and plateaued shortly after population isolation (Fig. 2). Plateaus positively correlated with admixture scenarios, with the highest PI observed for the highest admixture rate. PI did not considerably vary between admixture models, except during the time period separating the first from the second pulse of admixture. This result suggests that, under equal intensity of admixture (one pulse of 10 generations for SP and two pulses of 5 generations for MP), the number of admixture pulses could only be identified during generations separating them.

**Fig. 2.**
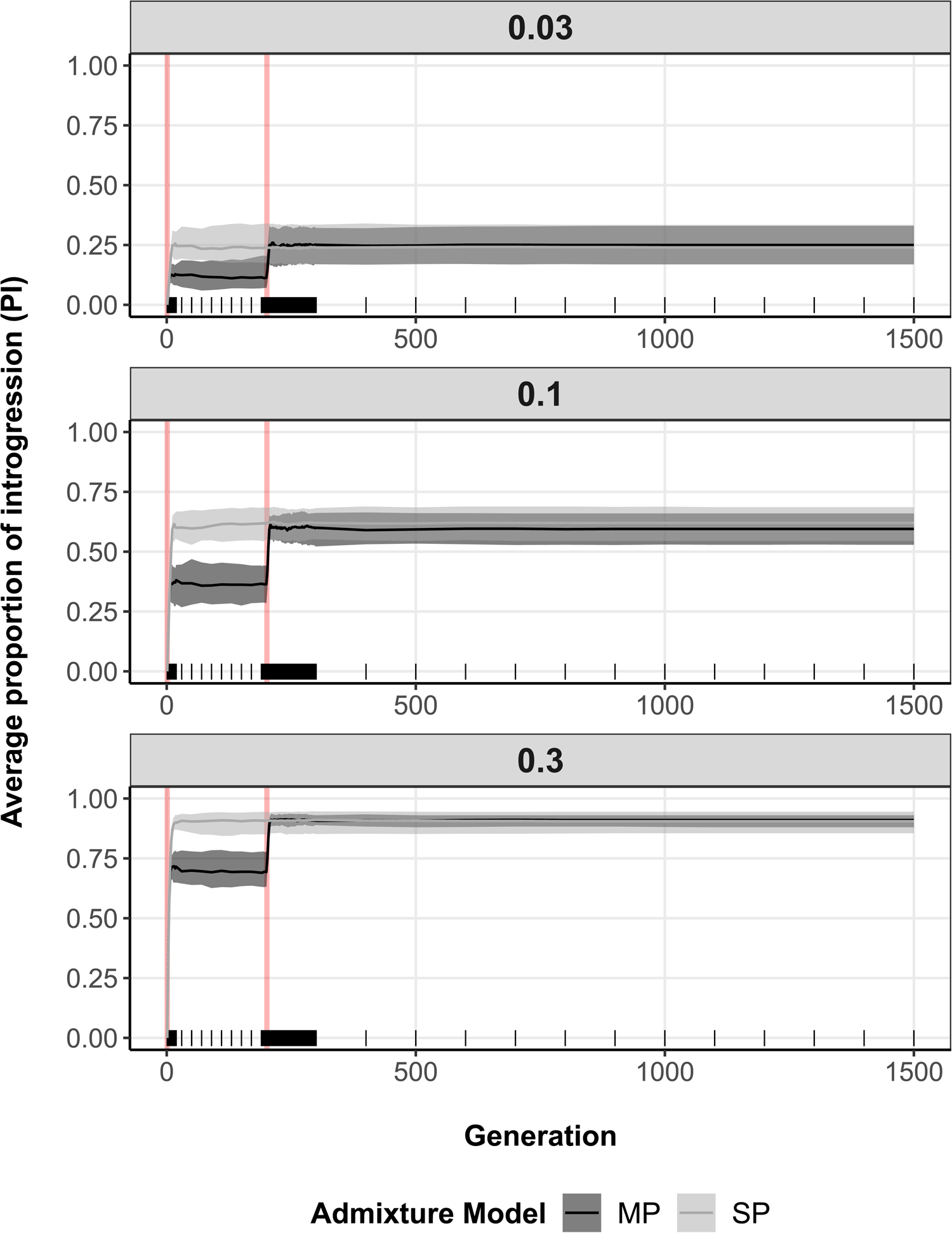
Average proportion of introgression (PI) (y axis) across generations (x axis), assuming a single pulse of admixture (gray) or multiple pulses of admixture (black). Shaded areas represent interquartile ranges delimited by the 25% (lower limit) and 75% (upper limit) quantiles of the distribution of introgression proportion averages (based on 100 simulation replicates) at each recorded generation (marked with tallies), while solid lines show median trends. Vertical red lines indicate the generation at which each admixture pulse starts (total time spent admixing: SP - generation 1 to 10, MP - generation 1 to 5 and 201 to 205). Results are separated by simulated admixture rates from source to sink population (0.03, 0.1, 0.3).

#### Lengths of introgressed sequences (LIS)

A general temporal pattern in LIS was observed across admixture models (Fig. 3). For the SP model, LIS was the longest at the beginning of the single admixture event, to shorten with advancing generations because of recombination. For the MP model, the same dynamics throughout generations applied, except it occurred twice, once after each admixture pulse. The peak in LIS associated with the second pulse of hybridization was considerably smaller than the one associated with the first admixture pulse for all admixture scenarios simulated. Average LIS associated with the second pulse of admixture shall always be smaller, as shorter fragments resulting from recombination occurring during the time interval separating admixture pulses will weigh down these averages. This is particularly evident for small admixture rates, where the input of longer sequences via gene flow is limited. Additionally, as time spent in isolation decreases between bouts of admixture, peaks will come closer and closer to each other until they can hardly or even no longer be teased apart (Appendix S1). This indicates that, to be successfully differentiated, admixture pulses need to be separated by a period of genomic isolation.

**Fig. 3.**
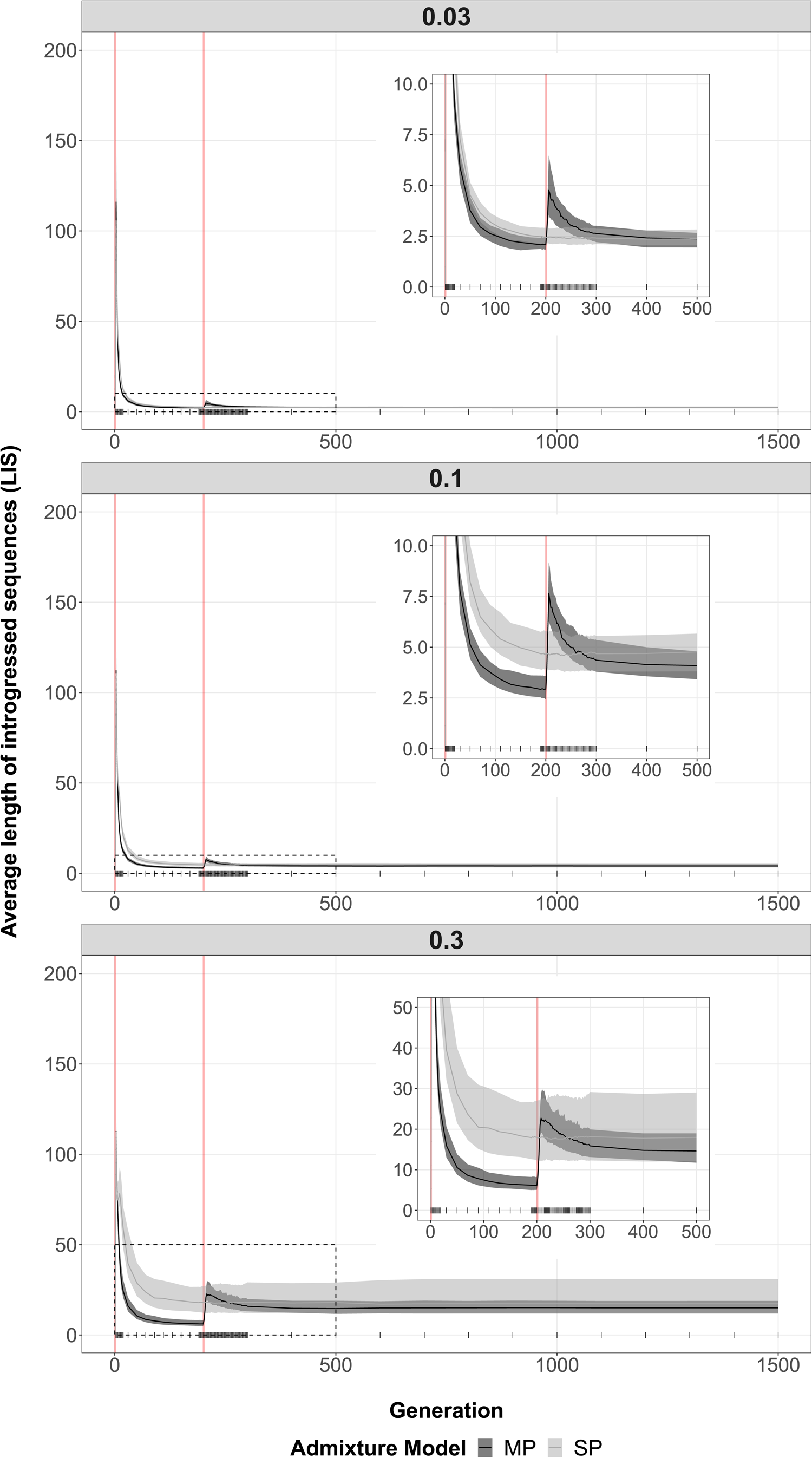
Average length of introgressed sequences (LIS) (y axis) across generations (x axis), assuming a single pulse of admixture (gray) or multiple pulses of admixture (black). Shaded areas represent interquartile ranges delimited by the 25% (lower limit) and 75% (upper limit) quantiles of the distribution of introgressed sequences’ length averages (based on 100 simulation replicates) at each recorded generation (marked with tallies), while solid lines show median trends. Vertical red lines indicate the generation at which each admixture pulse starts (total time spent admixing: SP - generation 1 to 10, MP - generation 1 to 5 and 201 to 205). Results are separated by simulated admixture rates from source to sink population (0.03, 0.1, 0.3). Insets are cropped sections of the full distribution indicated with a dashed rectangle.

Simulations also revealed that the speed at which introgressed fragments shorten differ among admixture scenarios as LIS declined faster for smaller admixture rate. In increasing order of admixture rates, decay rates between generation 3 and 200 were estimated at 0.190, 0.133, and 0.044 for SP and 0.283, 0.222, and 0.128 for MP, respectively. This dynamic may reflect a higher probability of homologous introgressed sequences to recombine together with increasing admixture rate, thus slowing down LIS decay. This same dynamic also contributed to the greater difference in LIS between SP and MP during generations separating hybridization pulses, when admixture increases. After the first pulse, there is more introgressed segments within the sink population under the SP model due to a longer period of hybridization (5 versus 10 generations of admixture for MP and SP, respectively), thus increasing the probability of two introgressed segments to recombine, ultimately lowering LIS decay.

Lastly, following the second pulse of admixture, average LIS stabilized for both SP and MP to subsequently plateau at similar values, values increasing with admixture rates simulated. Once again, the probability of two sequences originating from the source population to recombine within the sink population increases with higher admixture rates, resulting in higher LIS plateaus.

#### Intra-individual variation in introgressed sequences’ lengths (VIS)

Patterns of average VIS resembled closely those obtained for average LIS (Fig.3, Fig. 4). For both models (SP and MP) across admixture scenarios, the summary statistic increased at the beginning of hybridization pulses to decrease the following generations due to recombination homogenizing fragments’ sizes within individuals. This process occurred faster as gene flow between populations was small, with VIS decay rates between generation 3 to 200 estimated at 0.068, 0.053, and 0.030 for SP and 0.114, 0.109, and 0.072 for MP under admixture rates of 0.03, 0.1, and 0.3, respectively. The lower probability of two introgressed fragments to recombine for small admixture rates may explain the faster decline of VIS over time.

**Fig. 4.**
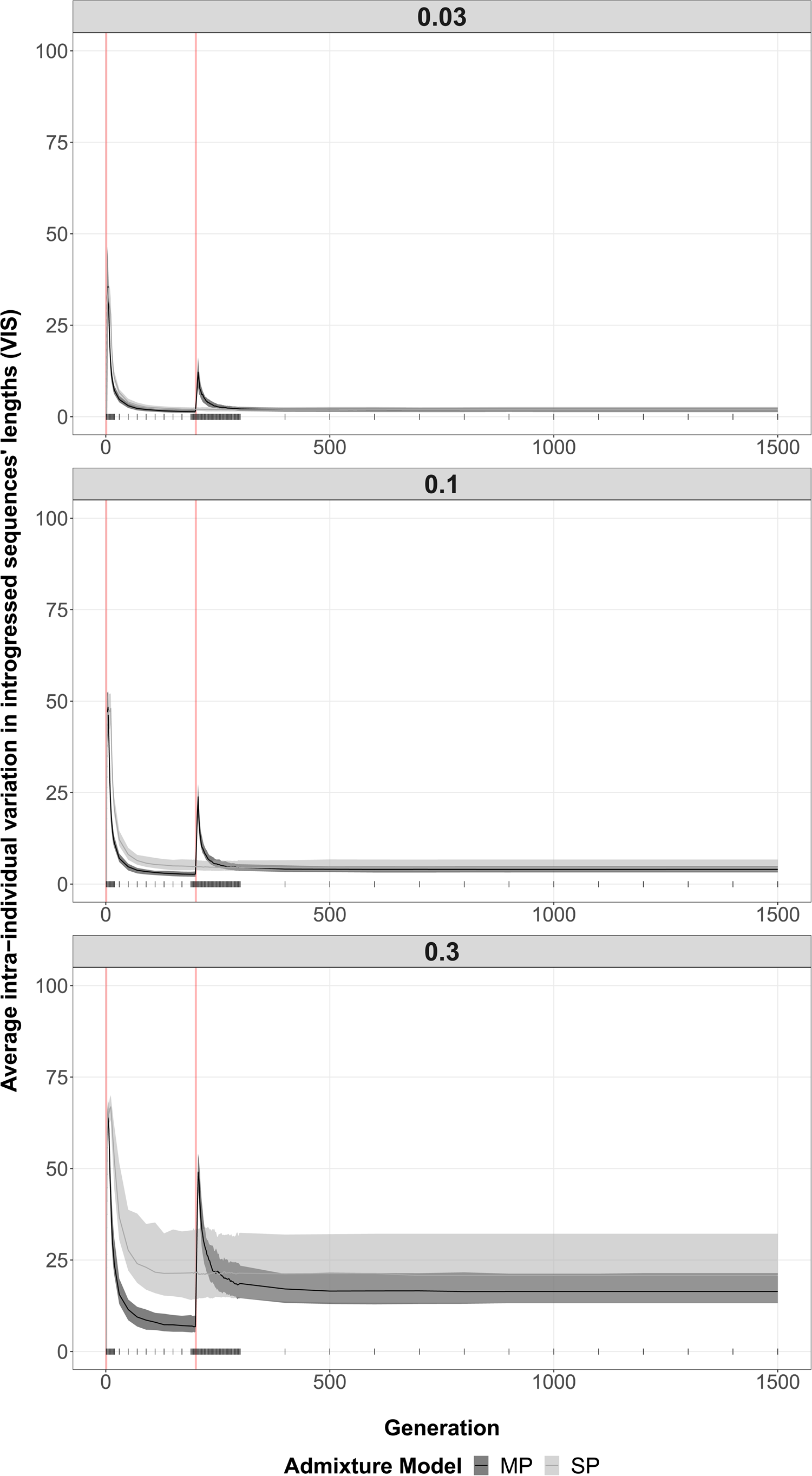
Average intra-individual variation in introgressed sequences’ lengths (VIS) (y axis) across generations (x axis), assuming a single pulse of admixture (gray) or multiple pulses of admixture (black). Shaded areas represent interquartile ranges delimited by the 25% (lower limit) and 75% (upper limit) quantiles of the distribution of variation averages (based on 100 simulation replicates) at each recorded generation (marked with tallies), while solid lines show median trends. Vertical red lines indicate the generation at which each admixture pulse starts (total time spent admixing: SP - generation 1 to 10, MP - generation 1 to 5 and 201 to 205). Results are separated by simulated admixture rates from source to sink population (0.03, 0.1, 0.3).

As observed for LIS, peaks of VIS associated with the second pulse of admixture were smaller than those characterizing the first pulse. Peaks’ heights were positively correlated with gene flow simulated, becoming taller as admixture rate increased. On the one hand, input of longer introgressed sequences from immigration, increasing the range of possible fragments’ lengths within individuals, may explain peaks observed when the second hybridization pulse occurred. Scenario-specific peaks’ heights, on the other hand, are the consequence of a higher expected frequency of recombination between introgressed segments for elevated admixture rates. Indeed, while recombination between introgressed and local segments mainly produces short fragments for small admixture rates, increased probability of recombination between two introgressed segments for higher admixture rates may produce a wider range of possible fragments’ lengths (i.e., long, intermediate, and short), overall increasing intra-individual variation. Note that, like observations made with LIS, generations of genomic isolation were necessary to produce admixture pulse-specific VIS peaks (Appendix S1).

Finally, as the last admixture pulse ended and the sink population remained genetically isolated, VIS decreased to ultimately stabilized to values increasing with admixture rate. Importantly, as observed previously with LIS, but to a lesser extent, a gap in VIS emerged between admixture models SP and MP during the time interval separating hybridization pulses as admixture rate increased. This phenomenon may result from the same combination of factors as those discussed above for LIS.

### Neanderthal introgression into Homo sapiens

To apply insights gained from simulations to our understanding of archaic introgression into modern humans, we analyzed *Homo neanderthalensis* and *Homo sapiens* genomes older than 10,000 years from the Allen Ancient DNA Resource (AADR) and summarized Neanderthal introgression using similar genomic statistics as those calculated on simulated data. While Neanderthal introgression was detected throughout ancient genomes over time (mean = 0.022, t_17_ = 28.47, P < 0.001), we observed no clear trends between PI and genomes’ age (Fig. 5a). All regressions performed, including linear, exponential, and GAM were non-significant (Table 1).

**Fig. 5.**
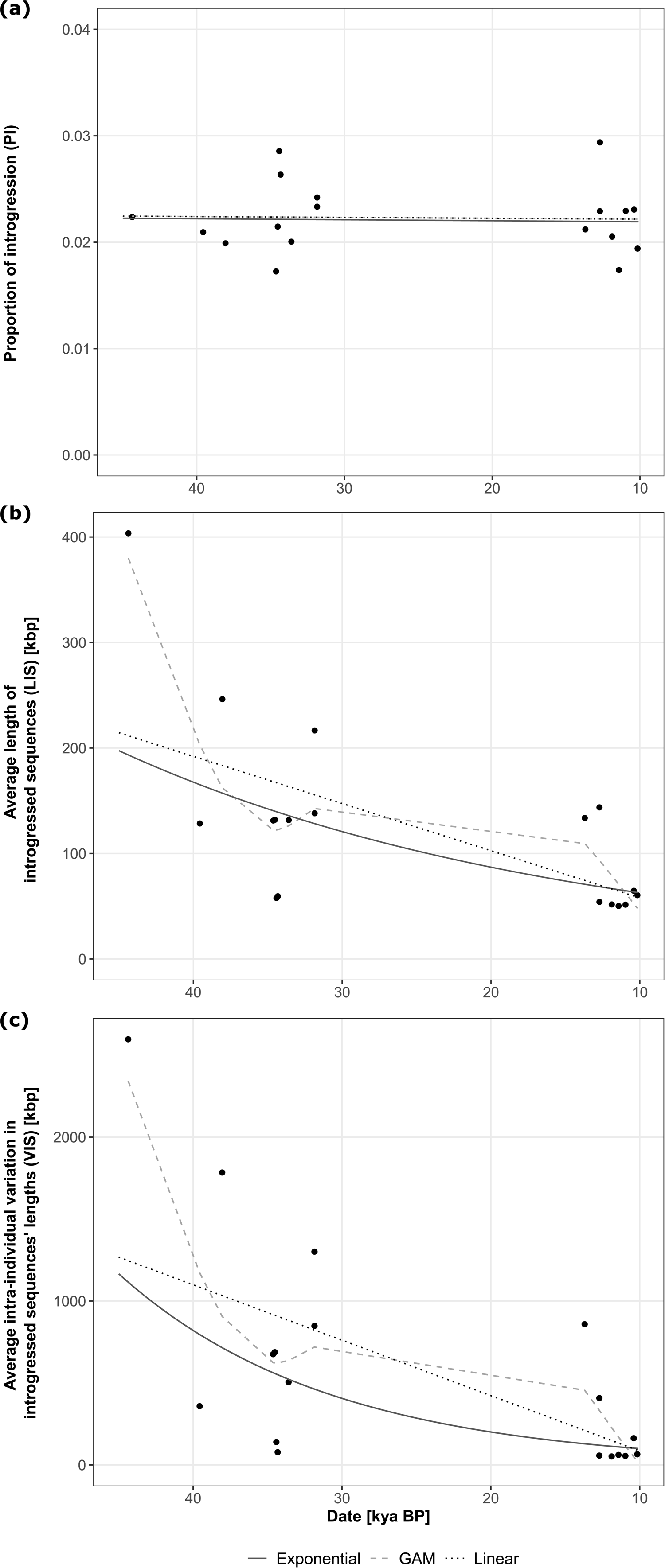
Genomic summary statistics (y axis) estimated for 18 ancient *Homo sapiens* genomes dating back from approximately 10 to 45 thousand years ago (x axis). (a) Proportion of introgression (PI). (b) Average length of introgressed sequences (LIS) [kbps]. (c) Average intra-individual variation in introgressed sequences’ lengths (VIS) [kbps]. The temporal dynamics of each parameter was evaluated using linear (dotted black line), log-linear/exponential (solid dark gray line), and GAM (dashed gray line) regressions.

**Table 1.**
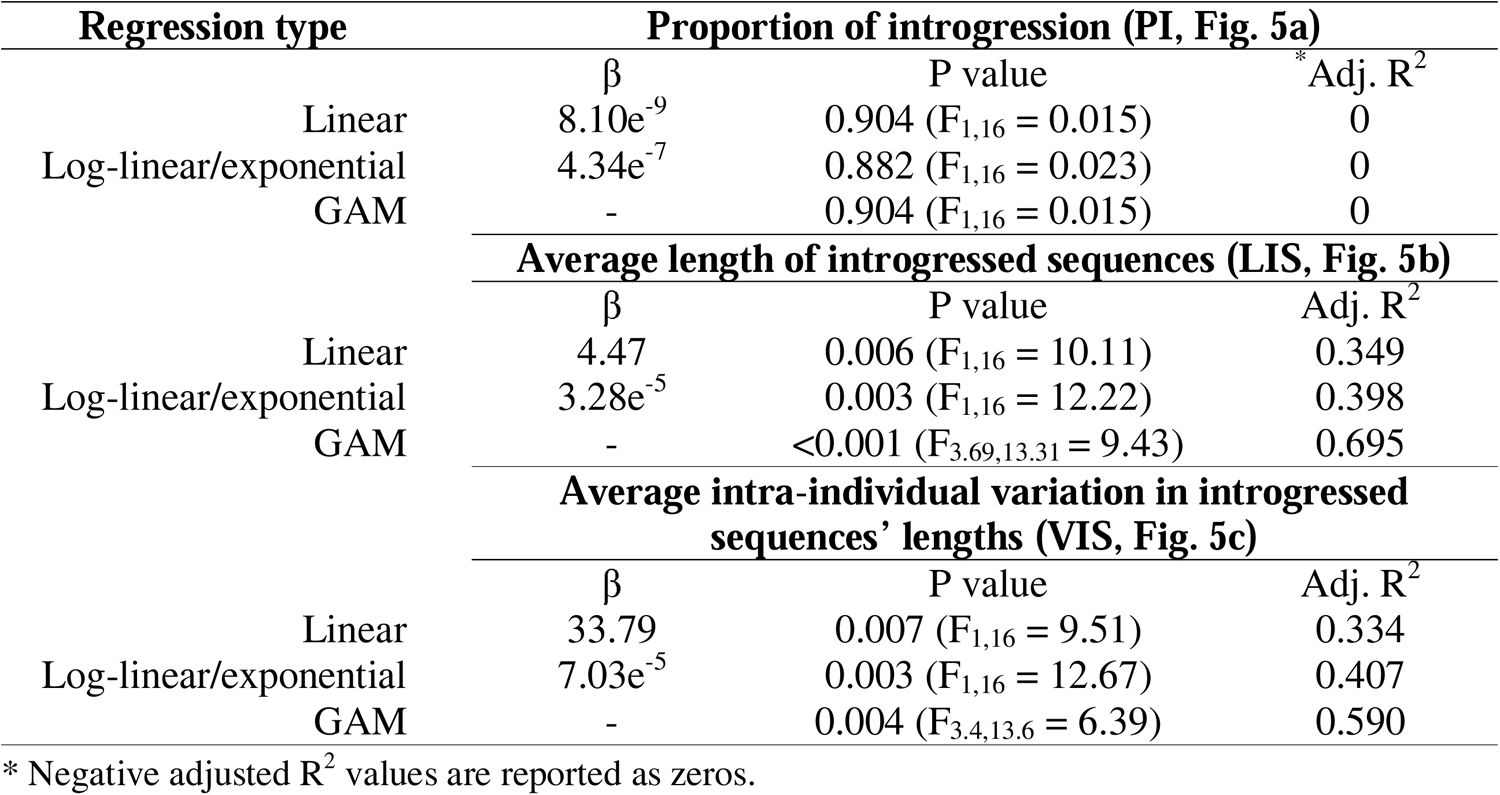
Statistics associated with all three types of regression performed (linear, log-linear/exponential, and GAM) between genomic parameters evaluated (proportion of introgression, PI; average length of introgressed sequences, LIS; and average intra-individual variation in introgressed sequences’ lengths, VIS) and age of ancient genomes retained for analysis. β: slope of regression line, Adj. R^2^: adjusted proportion of variation in genomic parameters explained by ancient genomes’ age.

With contrast to PI, both LIS and VIS (Fig. 5b, 5c) delineated a distinguishable pattern, increasing exponentially with genomes’ age. Indeed, while both linear and exponential regressions were significant (Table 1), the latter explained significantly more of the observed variation in LIS (χ_0_ = 429.08, P < 0.001) and VIS (χ_0_ = 476.15, P < 0.001) than the former. GAM regressions also were significant (Table 1) and provided similar insights with one additional noticeable feature. Between approximately 30 to 35 kya, another increase in LIS and VIS was observed, suggesting two possible pulses of admixture between archaic and modern human might have occurred. Nonetheless, we stress here that this additional peak is very speculative and may represent a statistical artifact resulting from the small sample size of available paleogenomes.

## Discussion

The growing field of paleogenomics offers the opportunity to explore novel empirical means to evaluate inter-lineage admixture and to improve our understanding of the complex evolutionary history of hybridizing species, including our own. Here, we used individual-based modelling to assess temporal patterns in three introgression summary statistics, including introgression proportions (PI), lengths of introgressed fragments (LIS), and intra-individual variation in introgressed fragments’ lengths (VIS) to determine whether they could be used to infer admixture pulses. Subsequently, we leveraged insights gained from simulations to evaluate whether targeted genomic statistics retain empirical value, estimating and interpreting them within the context of *Homo sapiens* and *Homo neanderthalensis* admixture history.

### Introgression proportions provide limited insights into lineage admixture history

Our results show that when one aims to infer the number of admixture pulses, this statistic may have limited value. Regardless of the admixture rate permitted between populations, our simulations demonstrated that a given amount of introgressive hybridization, occurring in either one (SP) or two (MP) pulses of admixture, largely resulted in similar PI. It is only during the inter-pulse period that SP and MP model could be differentiated from one another, suggesting that any temporal sampling failing to capture genomes from within this period would miss the signature of multiple admixture events. While PI is a widely used summary statistics in hybridization studies^12,32, 38–40^, this finding highlights the challenges associated with using PI to differentiate admixture pulses and supports the use of alternative statistics when trying to reconstruct inter-lineage hybridization history. For instance, many studies have investigated the admixture history between *Homo sapiens* and archaic hominins^19,22,35,41^, yet summary statistics like FFS (Fragment Frequency Spectrum), SFS (Site Frequency Spectrum), D-statistic, or segments of archaic origin inferred using a HMM (Hidden Markov Model) have been preferred over PI (but see^37^).

### Lengths of introgressed sequences and its variance within genomes could confidently distinguish admixture pulses

We showed that LIS and its variation within individuals (VIS) peaked during a short period of time around admixture pulses before decreasing because of recombination. These peaks, more pronounced for VIS, thus provide a means to approximate the timing and number of admixture events between two interbreeding lineages. Nonetheless, this also indicates that the time window (restricted here to the generation of admixture and the few dozen following it) to detecting a hybridization event is narrow. Importantly, simulations also revealed that this characteristic of introgressed fragments’ length statistics fades with weaker reproductive barriers between lineages (i.e., higher admixture rates), especially for LIS. Consequently, the strategy to detect multiple pulses of admixture appears to be context dependent and while priority should be given to samples spanning predicted times of admixture, the expected rate of reproductive isolation between interbreeding systems deserves consideration. While the ancient genome database for *Homo sapiens* may remain the largest to date^42^, and thus be used to test the empirical value of introgressed segments’ length statistics, ancient genomes are progressively being sequenced for non-human animals, plants, as well as pathogens and microorganisms^42,43^. As more of these genomes become available, our analytical framework could be used to study hybridization and differentiate between admixture pulses across a wider taxonomical breadth.

The term “admixture pulse” is commonly used in the literature, oftentimes described as a discrete period of time during which gene flow between distinct lineages is occurring^6,35,44,45^. However, this definition of an admixture pulse may be considered relatively broad. With our simulations, we found that multiple bouts of admixture are translated into peaks in introgressed sequences’ length statistics, as long as a period of genomic isolation separate hybridization events. Based on these results, we would like to propose a revisited definition of an admixture pulse as follow: “*a continuous series of generations during which genetic material is exchanged between biological entities, followed by a consecutive number of generations spent in genetic isolation*”. We believe this definition remains inclusive enough so it can be applied across a wide variety of hybridizing lineages, while being specifically associated with a quantifiable signal in readily computable introgression summary statistics.

### A clear admixture pulse detected prior to 45 kya between modern human and Neanderthal

Temporal patterns of introgression statistics estimated for ancient *Homo sapiens* genomes within the context of its admixture with Neanderthal supported the practicality of introgressed fragments’ length statistics in detecting pulse of hybridization. Regression analyses identified an increase in average LIS and VIS between approximately 33 to 45 kya, confirming our ancestors likely admixed with Neanderthal prior to 45 kya. Indeed, this estimate agrees with previous findings, placing the admixture time between 40 to 60 kya^6,46–49^. In addition, GAM regressions suggested the possibility of another peak, and thus another admixture event, between approximately 30 to 33 kya. However, a statistical artefact could not be excluded given the low number of paleogenomes post-dating this time period. The lack of temporal patterns found in introgression proportion further support this conclusion. Indeed, simulations demonstrated that under a multiple-pulse scenario, a stairs-like pattern in PI would be expected over time, in which plateaus reached increase in a stepwise fashion after each admixture pulse, a feature absent from ancient *Homo sapiens* genomes analyzed here. Finally, with a Neanderthal extinction time estimated between 35 to 50 kya^50,51^, a late admixture event between 30 to 33 kya remains unlikely. Addressing this uncertainty will ultimately necessitate additional paleogenomes spanning the time period from 15 to 40 kya to be sequenced.

### Limitations

While our study provides insightful observations on the temporal dynamics of three introgression summary statistics when distinct lineages admix on one or two discrete instances, our simulation framework could be easily expanded to simulate more than two admixture pulses. For both the LIS and VIS, we observed a noticeable decline in peak’s height associated with the second event of admixture. If this dynamic holds for subsequent hybridization pulses, there might be a limitation in the number of admixture events that could be detected using these statistics. Peaks would become shorter with any additional inter-breeding events until no more increase can be observed. Alternatively, it might be possible that peaks would decrease until reaching an equilibrium, where any additional admixture pulse would leave a weak, yet perceptible, genomic hybridization signal. Future studies simulating additional hybridization events using a similar framework to the one presented here would help shed some light on temporal patterns in introgression statistics expected when lineages admix on multiple occasions.

Particularly interested in patterns expected under neutrality, we did not consider the impact of selective forces in our simulations. Nevertheless, natural selection can impact the length distribution of introgressed segments and acknowledging this additional force may be important in some systems, including modern humans, where both negative and positive selection are hypothesized to have had a significant impact on the level of Neanderthal ancestry^4,32,33, 52–54^. Future work incorporating natural selection into the simulation framework may be performed to evaluate its impact on temporal patterns of length summary statistics. In addition, we did not incorporate gene flow with other populations into our simulations, which could impact the proportion of introgressed segments in the sink population^40,55^. Finally, while introgression and admixed segments’ lengths vary temporally, they also vary spatially^38,56,57^. To complement the temporal assessment conducted here, future research simulating introgression statistics using a spatiotemporal model would allow expected patterns to be described for more complex, realistic evolutionary scenarios.

## Conclusion

Our study demonstrates that, in addition to the well-established timing of hybridization^6,8,9,21,22^, length of introgressed sequences may be used to infer the number of admixture pulses between inter-breeding entities when assessed temporally. We also showed that a derived statistic, within-genome variance in introgressed fragments’ lengths, provides a valuable means to estimating the timing and number of hybridization pulses, particularly when admixture between lineages is limited. While simulation experiments featured signals of introgression bouts in the form of length and variation peaks, empirical estimation of these parameters on ancient *Homo sapiens* genomes accurately retrieved previously hypothesized hybridization events with Neanderthal. In summary, although this framework necessitates temporally distributed genomic data, it illustrates the practicality and limitations of a set of readily quantifiable introgression summary statistics to studying potentially intricate hybridization scenarios in an era where paleogenomics is popularizing.

## Methods

### Individual-based genomic simulations

#### Overview of the model

To evaluate genomics patterns emerging following temporal pulses of introgressive admixture between genetically differentiated lineages, we used a modified version of the *evolve()* function implemented within the R package “glads”^58^ (https://github.com/eriqande/glads) to conduct individual-based, forward-in-time, genomic simulations (see below for details on edits made to the function). The purpose of the individual-based model (IBM) implemented within this package is to explore how a range of genetic and demographic processes can influence genomic patterns between divergent lineages.

The IBM approach at the starting generation comprises individuals characterized by their genomic background (modeled as a single pair of homologous chromosomes), sexual identity (male or female), and their assigned population identity. Newborns have the same defining characteristics and will form the next generation of individuals within a population following reproduction. The pair of homologous chromosomes forming the genetic background of individuals is modeled using a suite of fixed characteristics: the length of homologs (in Mbps), the number and location of genetic markers (e.g., single-nucleotide polymorphisms - SNPs), the genotype (bi- or multi-allelic) at each genetic variant, the mutation rate per base pair, and the chromosome-wide average recombination rate per base pair. Values defined for each genomic parameter are shared among all individuals.

While various processes reflecting different evolutionary scenarios can be simulated using “glads”, we used the scenario simulating neutral evolution of chromosomes, including stochastic changes in population size through time (type = “dynamic”). For this scenario, first-generation individuals are created within a user-defined number of populations using the set of genomic and demographic parameters described above. Sexes are assigned to individuals using a binomial distribution with the probability of being a male equal to a given sex ratio. Then, reproduction within populations occurs by randomly forming mating pairs (consisting of one male and one female), generating haploid parental gametes based on recombination and mutation rates provided, and segregating haplotypes within mating pairs to produce the genotypes of newborns. Sexes of newborns are assigned as described above for first-generation individuals. The number of newborns produced per breeding pair is determined by two additional parameters; the population-specific mean number of descendants per breeding pair and the density-dependent demographic effect (to prevent exponential growth of populations). Ultimately, the number of offspring each breeding pair can have for a given generation will be calculated as: *O(t)* = *Pois(λ)* - *N(t)δ* (eq. 1), where *O(t)* is the number of offspring produced by a breeding pair at generation *t*, *λ* represents the population-specific mean number of descendants per breeding pair, *N(t)* represents the population size at generation *t*, and *δ* represents the density-dependent demographic effect (for details, see^58^). Following reproduction, newborns replace parental individuals and, if migration rates are specified for this generation, movement of offspring among populations occurs with probability of each newborn to migrate from population *i* to population *j* equal to migration rate *m_ij_*. Importantly, because migrants entering a new population are considered as potential mates during reproduction occurring in the next generation, this framework simulates inter-population gene flow. Eventually, the size of any population *i* in the next generation will be calculated as: 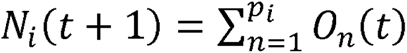 - *m_ij_N_i_(t)* + *m_ji_N_j_(t)*, where 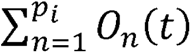 represents the overall reproductive output of all mating pairs within population *i* (*p_i_*) at generation *t* (see eq. 1), *m_ij_N_i_(t)* represents the number of individuals lost to emigration to population *j* at generation *t*, and *m_ji_N_j_(t)* represents the number of individuals gained from immigration from population *j* at generation *t*. The simulation ends when populations evolved for a provided number of generations *t* and genotypes of individuals at all SNPs within each population are returned as an output.

To increase flexibility of the IBM framework, we modified the *evolve()* function by incorporating the possibility to vary migration rates over time (previously fixed throughout generations), as well as to output genotypes of individuals within each population for a number of user-defined generations (previously only the genetic background of populations in the last generation was generated).

#### Initialization

We used a simulation framework considering the evolution of individuals distributed in two populations (a source - provider of migrants - and a sink - receiver of migrants - population). Both source and sink populations had an initial size of 100 individuals and evolved neutrally, this framework assuming no selection, equal sex ratio, and equal fitness of genotypes. Note, however, that demographic stochasticity was introduced in populations according to eq.1, so while fitness of genotypes was equal, the fitness of individuals within populations was not. Nonetheless, differences in among-individual fitness were stochastic and thus did not violate the neutrality assumption. The parameters *λ* = 4 and *δ* = 0.02 were identical between populations and were selected (1) based on fitness estimates for ancient hunter-gatherer populations^59^ and (2) their propensity to stabilize population sizes around the initial value of 100 individuals across generations, therefore avoiding the simulation of strong bottlenecks and population growths (see Appendix S2 and below for relevance). Populations were allowed to evolve for 1500 generations, providing a biologically relevant timeframe for simulation. Indeed, this corresponds grossly to the timing of admixture between Neanderthal and *Homo sapiens* during their OOA (Out-Of-Africa) expansion (∼1500-1800 generations ago^6^).

Finally, to realistically mimic the genomic dynamics of a biological system in our simulations, we again leveraged the literature available for *Homo sapiens* to choose appropriate genetic parameters. Following Kong et al.^60^, the genomic background of each individual was modeled as a single pair of homologous chromosomes with an average recombination rate of 1.2e^-8^ per base pair. Each chromosome had a length of 150 Mbp with 200 evenly spaced genetic markers. The major objective of these simulations being to track the fate of introgressed sequences through time and not to reproduce faithful population genetic diversity, mutation rate was set to 0 and source and sink populations at generation 1 were fixed for alternate alleles at all 200 loci (allele 1 for sink and 2 for source). This way, the introgressed allele could always be differentiated from the local allele, facilitating downstream analyses.

#### Simulations

To assess whether differences in temporal patterns of introgression exist between genomes experiencing single or multiple pulses of introgressive hybridization, two contrasting admixture models were considered: one simulating a single pulse of admixture (SP model), and another considering two independent pulses of admixture (MP model). Both models shared general IBM parameters described previously but differed in how hybridization was simulated. For the SP model, admixture only happens once from generation 1 to generation 10, followed by 1490 generations of genetic isolation and neutral evolution (Fig. 1a). Contrastingly, for the MP model, admixture happens twice, once from generation 1 to generation 5, and once from generation 201 to generation 205 (Fig. 1b). Admixing time was kept identical between models so that the effect of the number of pulses could be disentangled from the effect of admixture intensity on genomic summary statistics investigated (see below). To represent the effect of admixture, migration was set as unidirectional in both SP and MP models, always from the source to the sink population, to keep the former population free of the latter population’s allele. This way, we avoided dealing with uncertainties pertaining to shared ancestries between the source and sink populations in estimating introgression summary statistics.

In addition to the number of hybridization pulses, we also investigated the impact admixture rate may have had on the temporal dynamics of genomic summary statistics (note that here what we call admixture rate is in fact the migration rate (*m_ij_*) between populations in “glads”). SP and MP models were each run three times using a different admixture rate (referred to as scenarios). Specifically, SP and MP models were simulated with an admixture rate from source to sink population of 0.03, 0.1, and 0.3. In summary, three scenarios were considered and implemented for each admixture model (SP and MP), scenarios differing only by the intensity of interpopulation admixture simulated. We mentioned earlier those parameters incorporating demographic stochasticity into simulations (*λ* and *δ* respectively) were chosen as they allowed sizes of source and sink populations to stabilize around the initial population size provided of 100 individuals over time (Appendix S2), allowing temporal patterns of introgression to reflect mainly the impact of gene flow levels, as opposed to the impact of population size variation. Finally, variability in genomic patterns of introgression stemming from the stochastic evolution of chromosome pairs within the sink population was accounted for by simulating each scenario per admixture model 100 times. Genotypes of all individuals within sink and source populations were recorded for select timepoints to estimate introgression summary statistics of interest and their evolution over time. Particularly, simulation outputs were generated every 2 generations from generation 1 to 20, every 20 generations from generation 30 to 190, every 2 generations from generation 192 to 300, and every 100 generation from generation 400 to 1500. Consequently, we have a more detailed view of hybridization periods relative to isolation periods.

#### Estimation of population-level genomic summary statistics

Using simulation outputs of recorded generations, we estimated three summary statistics for sink population replicates to describe genomic patterns of introgression. These include (1) the average proportion of introgression (PI), (2) the average length of introgressed sequences (LIS), and (3) the average intra-individual variation in the lengths of introgressed sequences (VIS). The proportion of introgression was calculated for each individual as the number of introgressed alleles across homologs, divided by the total number of haploid markers (400). Individual-specific proportions of introgression were subsequently averaged across all individuals forming the sink population. To estimate the lengths of introgressed sequences, each homologous chromosome present within individuals assigned to the sink population was scanned for continuous tracts of the source population’s allele (allele 2). This was performed by counting the number of consecutive introgressed alleles, whenever at least one was encountered along the sequence of a homolog, and considering the suite of such alleles as an introgressed segment. Ultimately, the population-level estimate for the lengths of introgressed sequences was estimated by averaging the number of consecutive introgressed alleles forming a segment over all inferred segments across individuals. Because all 200 loci simulated were evenly spaced along a chromosome, segment length measured in number of loci or bp is proportional. For simplicity, we only estimated and presented the former. Lastly, intra-individual variation in introgressed sequences’ lengths was estimated first by calculating the standard deviation of all introgressed segments inferred for each sink individual separately, and then by averaging these values across all individuals within that population. Note that for each recorded generation, genotypes of immigrants and offspring born from the reproduction between two immigrants (pure source individuals genetically) were not considered in the estimation of these summary statistics, as they cannot be regarded as introgressed. Functions needed to conduct simulations and compute all three genomic summary statistics were compiled into a R package, dependent on “glads”, entitled “companions4glads” provided as Supplementary Material.

Temporal changes in genomic summary statistics were eventually assessed by evaluating the distribution of population-level averages estimated for all 100 simulation replicates across recorded generations. The estimation of the decay rate of LIS and VIS over time was performed for all admixture rates separately by regressing median values of population-level averages across generations using R *nls()* function and a negative exponential equation: *y(t)* = *y_f_* + *(y_0_ - y_f_)e^-at^* + *ɛ*, where *y(t)* is the value of the summary statistic at generation *t, y_0_* is the value from which the summary statistic starts to decay, *y_f_* is the value toward which the summary statistic decays, *ɛ* is the rate of decay, and *ɛ* the residual error. All analyses were conducted in R version 4.2.2^61^ and 4.2.3^62^.

### Ancient genome analysis

#### Data collection and filtering

To determine whether patterns of introgression observed in simulations could be replicated with empirical data, analyses like those conducted on simulated data were performed using ancient genomes available from the Allen Ancient DNA Resource (AADR) (<u>https://</u> <u>reich.hms.harvard.edu/allen-ancient-dna-resource-aadr-downloadable-genotypes-present-day-</u> <u>and-ancient-dna-data</u>). The whole database (version v50.0, release 10/10/2021) was downloaded and subsequently filtered in R. Primarily focusing on Neanderthal introgression into the genome of *Homo sapiens*, we selected two distinct Neanderthal genomes with the highest coverage, namely the Altai_published.DG and Vindija_snapAD.DG genomes. Among all *Homo sapiens* genomes, we kept only those with a base coverage greater than four and removed duplicated samples, retaining genomes with the highest number of SNP calls on autosomes. These filters were applied (1) to keep genomes optimizing the accuracy of inferences while minimizing genome loss for adequate temporal sampling and (2) to ensure genome independence. Finally, one outgroup genome (chimpanzee – Chimp.REF) was selected for computation of f4 ratios (see below). Overall, 935 genomes were retained and used for downstream analyses, including 932 *Homo sapiens* genomes, two Neanderthal genomes, and one outgroup (chimpanzee) genome.

#### Estimation of genomic summary statistics

Genome-wide average of Neanderthal introgression proportion was estimated for 931 of the 932 *Homo sapiens* individuals using the R package “admixr”^63^. The missing genome was the contemporary African genome A_Dinka-4.DG we used as a non-introgressed reference. Proportions of introgressions (PIs) were measured as f4 ratios^64^ (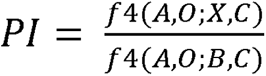, eq. 2), a statistic capable of inferring ancestry fractions in admixed populations given genotypes from a presumed gene donor population (i.e., Altai_published.DG; B in eq. 2), genotypes from a population closely related to the gene donor (i.e., Vindija_snapAD.DG; A in eq. 2), genotypes from putatively introgressed populations (i.e., all 931 contemporary and ancient *Homo sapiens* genomes; X in eq. 2), genotypes from a non-introgressed reference population (i.e., A_Dinka-4.DG; C in eq. 2), and genotypes from an outgroup species (i.e., Chimp.REF; O in eq. 2).

The number, location, and length of Neanderthal introgressed segments were assessed for the same 931 *Homo sapiens* genomes using the software “IBDmix”^34^. The analysis was performed using genotypes called from the Altai Neanderthal (Altai_published.DG) as the archaic reference. End positions of introgressed segments were reported as inclusive (-i) so that the one reported is the last increasing log odds, as opposed to the first one decreasing log odds. All remaining parameters were kept to default.

#### Statistical analyses

Interested in temporal patterns of introgressions, we selected among all 931 *Homo sapiens* genomes analyzed only those from Eurasia and the Americas that were 10,000 years old or older. Neanderthals and *Homo sapiens* likely temporally overlapped from when modern humans started to disperse outside Africa (∼75 kya)^65^ and into Eurasia (∼47-55 kya)^48,66^ and Neanderthal extinction (∼35-50 kya)^50,51^. Consequently, we believe 10,000 years old and older genomes should provide an appropriate time window to study consequences of introgressive hybridization between the two hominin species. In total, we retained 18 ancient human genomes with significant genome-wide proportions of Neanderthal introgression (Z score > 3) for statistical analysis (see Appendix S3 for details). For each individual genome, “IBDmix” reported the start and end position of introgressed segments (log odds ≥ 3) per chromosome. More interested in genome-wide rather than chromosome-specific patterns of admixture, we averaged the lengths of introgressed segments per chromosome, and then across chromosomes, generating a single mean value per individual (LIS). Intra-individual variance in introgressed sequences’ lengths (VIS) was similarly processed. Standard deviation of segments’ lengths was calculated for each chromosome independently and averaged across chromosomes, providing a single mean value per individual.

Temporal patterns of Neanderthal introgression into the genome of modern humans were evaluated using a set of three distinct regression analyses, including a linear, exponential (log-linear), and GAM regression. Each genomic summary statistic (response) was regressed over the average age of each sample (predictor), estimated using radiocarbon dating and recorded within the AADR. We chose exponential (log-linear) regression as we observed with genomic simulations an exponential relationship between introgressed fragment summary statistics (including both LIS and VIS) and time (see Results). Nonetheless, as ancient genomes available are unlikely to be as well temporally distributed as generation snapshots taken with simulations, we also decided to test for a linear relationship between these variables. While the scarcity of ancient genomes available may hinder an exponential relationship, a linear interconnection between introgressed segment summary statistics and passing generations may still be detectable and provide some insights. To evaluate which of the linear or exponential regressions explained the most variation in genomic summary statistics, we conducted a likelihood ratio test as implemented in the function *lrtest()* from the “lmtest”^67^ R package.

Finally, the regression assuming a generalized additive model (or GAM) was performed in R using the “mgcv”^68^ package. This method was selected as it may be particularly useful when trying to evaluate a subtler relationship between two variables. Indeed, the non-linear smoothing of the relationship between covariates may allow more complex trends hidden with simpler regression methods to be put forward. Note, however, that the shape of the relationship may largely be influenced by the basis dimension used to represent the smoothing term (also known as the *k* parameter). To find the *k* value fitting the data best (the *k* value leaving no more observable patterns in the residuals), we used the function *k.check()* to identify, over all possible *k* values, the one with the highest p value (the higher the p value, the more likely it is that no more patterns exist in the residuals). When ties for the highest p-value were observed, we chose the lowest *k* value resulting in this p-value. This selection process was repeated for every GAM regression involving a different genomic summary statistic.

## Data availability

No novel datasets were generated as part of this study. The whole set of functions used to conduct simulations and estimate introgression summary statistics were compiled into a small R package complementary to “glads” (https://github.com/eriqande/glads), entitled “companions4glads”, available as Supplementary Material (Appendix S4). Lastly, details on genomes used to generate Fig. 5 are presented in Appendix S3.

## Acknowledgments

We thank Stephan Weber and Jose Manuel Nunes for their support with the use of the high-performance cluster of the AGP lab at the University of Geneva. This research was supported by grants from the Swiss National Science Foundation (Grant number: 31003A_182577) to M.C and (Grant number: P5R5PB_203169) to C.S.Q.

## Author contributions

**L.N.D.S** designed the study, performed the simulations, analyzed ancient genomes, interpreted the results, and led the writing of the manuscript. **C.S.Q** and **M.C** designed the study, interpreted the results, and wrote the manuscript.

