## Appendix for "Temporal variation in introgressed segments’ length statistics sheds light on past admixture pulses"

### Appendix S1. Extra simulations (referred to as simulation A, B, C, D, E, and F; see Table S1 for details) evaluating the impact of the time of admixture pulse on temporal patterns of introgressed sequences’ length statistics LIS and VIS (Figure S1, Figure S2). Simulations were performed using identical IBD parameters as those described in the main text. Only exceptions are (1) the time at which the second pulse of admixture occurred (Table S1), (2) the total number of generations simulated (250 instead of 1500), and (3) generation snapshots recorded (all generation from 1 to 250 instead of intervals between generation 1 to 1500).

#### Table S1. Simulation parameters. Listed are the identification given to each simulation performed, the generation at which each admixture pulse started and ended, and generations where populations’ genomic composition were recorded.

| **Simulation ID*** | **First admixture pulse** | | **Second admixture pulse** | | **Generations recorded** |
| --- | --- | --- | --- | --- | --- |
|  | **Start** | **End** | **Start** | **End** |  |
| A | 1 | 5 | 201 | 205 | 1 to 250 |
| B | 1 | 5 | 101 | 105 | 1 to 250 |
| C | 1 | 5 | 51 | 55 | 1 to 250 |
| D | 1 | 5 | 26 | 30 | 1 to 250 |
| E | 1 | 5 | 11 | 15 | 1 to 250 |
| F | 1 | 5 | 6 | 10 | 1 to 250 |

* Simulation A and F are in all aspect identical to MP and SP simulations, respectively, presented in Figure 3 and 4 of the main text, except that genomic evolution of the sink population was followed for 250 instead of 1500 generations.

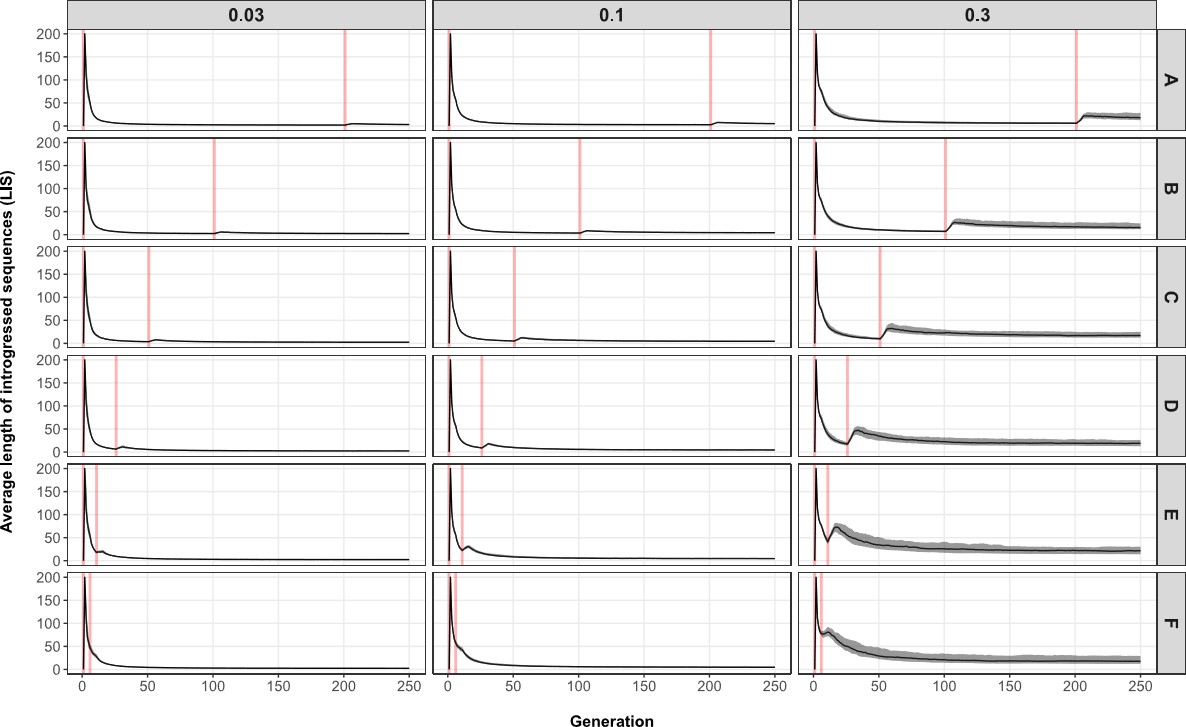

#### Figure S1. Average length of introgressed sequences - LIS - (y axis) across generations (x axis), separated by simulation ID (A-F; rows) and admixture rate permitted between source and sink populations (0.03, 0.1, 0.3; columns). Shaded areas represent interquartile ranges delimited by the 25% (lower limit) and 75% (upper limit) quantiles of the distribution of introgressed sequences’ length averages (based on 100 simulation replicates) at every generation, while solid lines show median trends. Vertical red lines indicate the generation at which each admixture pulse starts (total time spent admixing: 10 generations – see Table S1 for details).

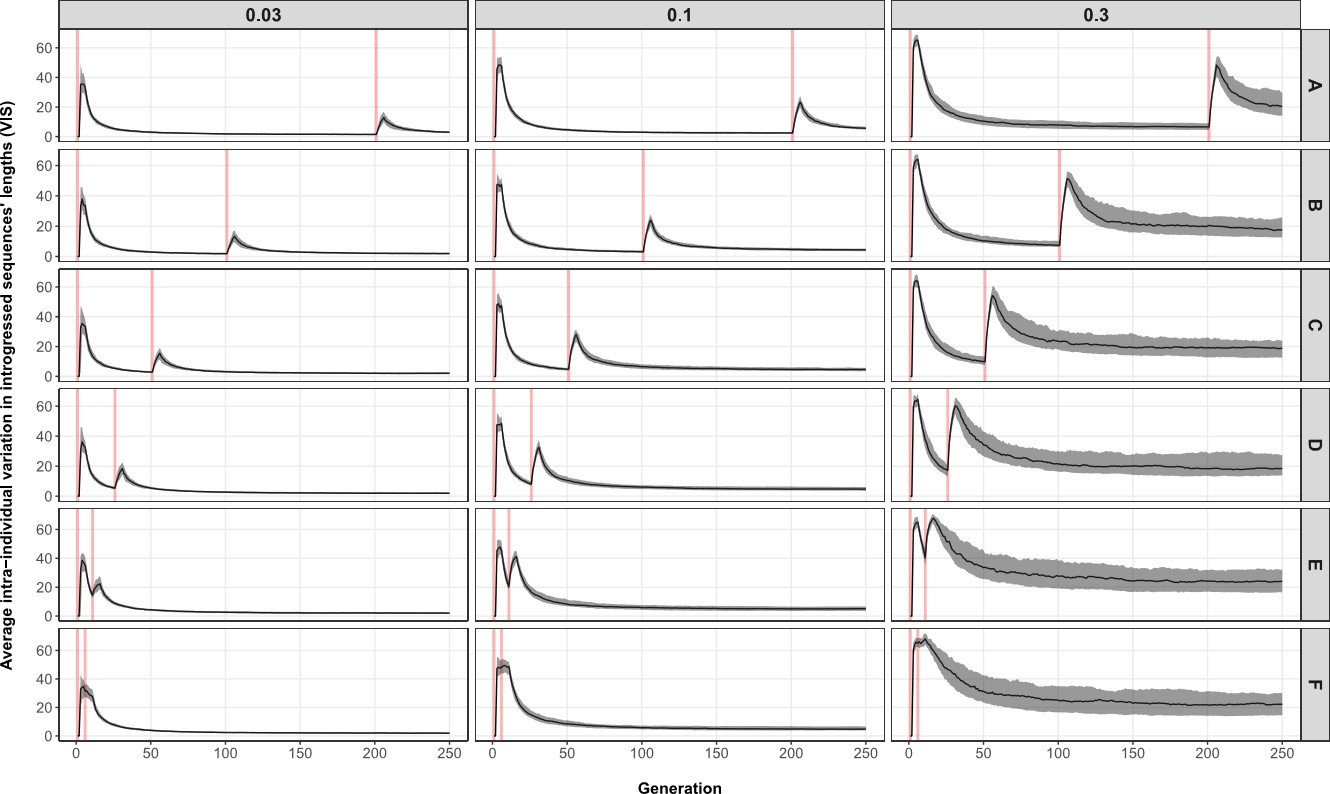

#### Figure S2. Average intra-individual variation in introgressed sequences’ lengths - VIS - (y axis) across generations (x axis), separated by simulation ID (A-F; rows) and admixture rate permitted between source and sink populations (0.03, 0.1, 0.3; columns). Shaded areas represent interquartile ranges delimited by the 25% (lower limit) and 75% (upper limit) quantiles of the distribution of variation averages (based on 100 simulation replicates) at every generation, while solid lines show median trends. Vertical red lines indicate the generation at which each admixture pulse starts (total time spent admixing: 10 generations – see Table S1 for details).

### Appendix S2. Population sizes averaged over all 100 simulation replicates (y axis) provided for recorded generations (x axis) under the single-pulse - SP - (a) and multiple-pulse - MP - (b) admixture models. Population size averages are separated by population identity (columns) and migration scenarios (rows). Shaded areas represent interquartile ranges delimited by the 25% (lower limit) and 75% (upper limit) quantiles of the distribution of population sizes for a given generation. Vertical red lines indicate the generation at which each admixture pulse starts (total time spent admixing: SP - generation 1 to 10, MP - generation 1 to 5 and 201 to 205).

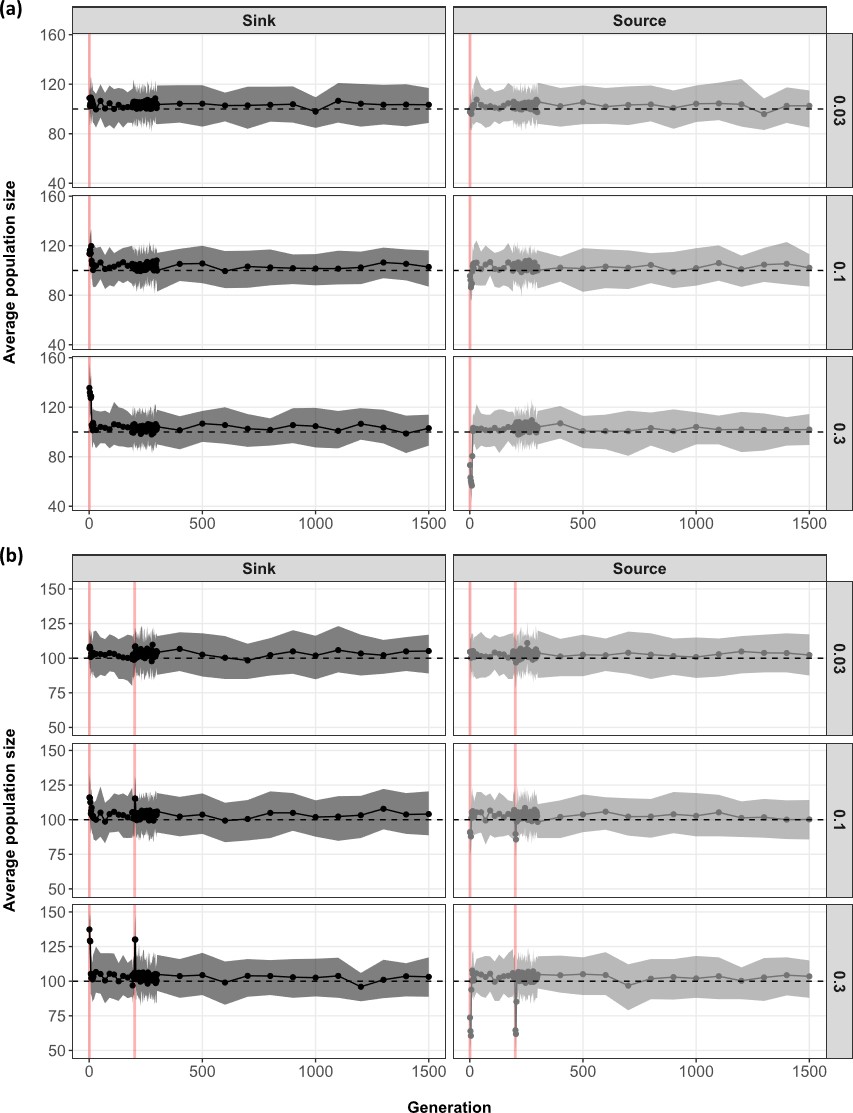

### Appendix S3. General information and summary statistics estimated for ancient genomes retained for analysis. AADR: Allen Ancient DNA Resource, LIS: average length of introgressed sequences, VIS: average intra-individual variation in introgressed sequences’ lengths.

| **General Information** | | | | | |  | **Genomic Summary Statistics** | | |
| --- | --- | --- | --- | --- | --- | --- | --- | --- | --- |
|  |  |  |  |  |  |  |  | **Sequence length** | |
| **Genome ID (AADR)** | **Category** | **Date (years before 1950)** | **Country** | **Lat** | **Long** |  | **Proportion of introgression** | **LIS (bp)** | **VIS (bp)** |
| Chimp.REF | Outgroup | 0 | - | - | - |  | - | - | - |
| Vindija_snpAD.DG^1^ | Neanderthal | 50,000 | Croatia | 46.30 | 16.07 |  | - | - | - |
| Altai_published.DG^2^ | Neanderthal | 110,450 | Russia | 51.41 | 84.69 |  | - | - | - |
| A_Dinka-4.DG^3^ | African | 0 | Sudan | 8.78 | 27.4 |  | - | - | - |
| I1954^4^ | non-African | 10,162 | Iran | 34.45 | 48.12 |  | 0.019 | 60,472.41 | 65,227.55 |
| Sumidouro5.SG^5^ | non-African | 10,403 | Brazil | -19.54 | -43.94 |  | 0.023 | 64,680.83 | 163,197.04 |
| AHUR_2064.SG^5^ | non-African | 10,962 | USA | 37.41 | -122.08 |  | 0.023 | 51,504.89 | 55,330.42 |
| USR1.SG^6^ | non-African | 11,425 | USA | 64.22 | -145.70 |  | 0.017 | 50,247.18 | 61,344.03 |
| I11974.SG^7^ | non-African | 11,885 | Chile | -31.92 | -71.50 |  | 0.021 | 51,716.53 | 51,954.19 |
| PES001.SG^8^ | non-African | 12,711 | Russia | 61.23 | 38.91 |  | 0.029 | 143,737.95 | 408,093.53 |
| Anzick_realigned.SG^9^ | non-African | 12,712 | USA | 45.99 | -110.66 |  | 0.023 | 54,125.26 | 57,232.40 |
| Bichon.SG^10^ | non-African | 13,698 | Switzerland | 47.10 | 6.87 |  | 0.021 | 133,693.52 | 858,544.38 |
| Yana_old.SG^11^ | non-African | 31,850 | Russia | 70.72 | 135.42 |  | 0.023 | 138,127.89 | 848,506.36 |
| Yana_old2.SG^11^ | non-African | 31,850 | Russia | 70.72 | 135.42 |  | 0.024 | 216,642.30 | 1,301,361.45 |
| NE20^12^ | non-African | 33,592 | China | 45.40 | 127.03 |  | 0.020 | 131,659.59 | 505,118.02 |
| Sunghir4.SG^13^ | non-African | 34,323 | Russia | 56.18 | 40.50 |  | 0.026 | 59,484.27 | 77,722.54 |
| PM1^14^ | non-African | 34,415 | Romania | 45.19 | 23.75 |  | 0.029 | 57,798.13 | 139,800.49 |
| Sunghir3.SG^13^ | non-African | 34,517 | Russia | 56.18 | 40.50 |  | 0.021 | 132,008.78 | 687,931.19 |
| Sunghir2.SG^13^ | non-African | 34,629 | Russia | 56.18 | 40.50 |  | 0.017 | 131,249.24 | 675,741.74 |
| Kostenki14^15^ | non-African | 38,052 | Russia | 51.23 | 39.30 |  | 0.020 | 246,293.66 | 1,784,189.68 |
| Tianyuan^16^ | non-African | 39,565 | China | 39.66 | 115.87 |  | 0.021 | 128,482.61 | 358,461.89 |
| Ust_Ishim_published.DG^17^ | non-African | 44,366 | Russia | 57.70 | 71.10 |  | 0.022 | 403,535.46 | 2,597,018.39 |
