## Appendix S4 for "Temporal variation in introgressed segments’ length statistics sheds light on past admixture pulses": companions4glads_1.2.pdf

### Package ‘companions4glads’

April 5, 2023

**Title** Analysis of Simulation Outputs from Glads

**Version** 1.2

**Description** 'companions4glads' is a package containing analytical functions for outputs generated by the R package glads. (https://github.com/eriqande/glads).

**License** GPL-2

**Imports** progress,
parallel,
HiddenMarkov,
foreach,
doParallel,
LaplacesDemon,
glads

**Encoding** UTF-8

**Roxygen** list(markdown = TRUE)

**RoxygenNote** 7.2.3

#### R topics documented:

**Index** 9

`compute_introgression_lengths`

*Compute Lengths of Introgressed Sequences*

#### Description

This function computes the lengths of introgressed sequences within a select population. Length is calculated for each individual and pair of homologous chromosome within individuals separately (but see arguments `pool` and `stats` for other outputs).

#### Usage

```
compute_introgression_lengths(
  x,
  pop = 1,
  allele = c(native = 1, introgressed = 2),
  loc.pos,
  stats = NULL,
  exclude = F,
  pool = F
)
```

#### Arguments

| | |
|---|---|
| `x` | Output from `evolve2.0` in `struct` format. |
| `pop` | *numeric* Which simulated population should the length of introgressed sequences be estimated on. |
| `allele` | *numeric* A vector describing (in order) the integer for the native and introgressed allele. |
| `loc.pos` | *numeric* A vector describing the position of each marker (same as the one used for simulation with `evolve2.0`). |
| `stats` | *character* One of "length" or "nb.loci". Specifying this argument triggers the estimation of intra-individual standard deviation in introgressed sequences length (in bp –> "length" or number of loci –> "nb.loci"). |
| `exclude` | *logical* Whether or not introgressed sequences of length = 1bp or 1 locus should be excluded (TRUE = yes, FALSE = no). |
| `pool` | *logical* Whether results should be pooled together (one vector with all lengths across all homologs and individuals). |

#### Note

Please note that if there is no introgression within the studied population, a series of 3 zeros (if `stats` = `NULL` and `pool` = `TRUE`) or zeros (for descriptive statistics and data) will be produced (if `stats` = `c("length", "nb.loci")`). If `stats` = `NULL` and `pool` = `FALSE`, an empty list is returned.

---

`compute_introgression_peaks`
*Compute Introgression Peaks (Islands or Valleys of Introgression)*

---

#### Description

This function takes the output of `evolve2.0` and assess peaks of high (island) and low (valley) introgression. Peaks are assessed based on posterior probabilities of states inferred using a hidden markov model and permutation test (3 states: Background, high, low). This function also computes the length of islands and valleys of introgression as well as the distance between them.

#### Usage

```
compute_introgression_peaks(
  x,
  pop = 1,
  introgressed.allele = 2,
  loc.pos,
  cores = 1,
  n.perm = 999,
  exclude = T
)
```

#### Arguments

| | |
|---|---|
| `x` | Output from `evolve2.0` or in `struct` format. |
| `pop` | *numeric* Which simulated population should the peaks be estimated on. |
| `introgressed.allele` | *numeric* The code for the introgressed allele. |
| `loc.pos` | *numeric* A vector describing the position of each marker (same as the one used for simulation with `evolve2.0`). |
| `cores` | *numeric* Number of cores to use for HMM analysis. |
| `n.perm` | *numeric* N number of permutations to assess significance of island and valleys of introgression inferred with HMM. |
| `exclude` | *logical* Whether island and valleys of length 1 should be excluded (TRUE) or not (FALSE). |

---

`compute_introgression_proportions`
*Compute Introgression Proportions*

---

#### Description

This function computes different summary statistics on simulation outputs (in `struct` format).These statistics include individual-specific introgression, average introgression, range of introgression, quantiles of the distribution of introgression and standard deviation of introgression.

##### Usage

```
compute_introgression_proportions(x, pop = 1, introgressed.allele = 2, loc.pos)
```

##### Arguments

| | |
|---|---|
| x | Output from evolve2.0 in struct format. |
| pop | *numeric* Which simulated population should the summary statistics be calculated on. |
| introgressed.allele | *numeric* The code for the introgressed allele. |
| loc.pos | *numeric* A vector describing the position of each marker (same as the one used for simulation with evolve2.0). |

##### Value

$Prop.introgression.table: A list with individual-specific introgression proportions.

$mean.introgression: Chromosome-wide introgression (average across individuals).

$range.introgression: Range of introgression proportions across individuals.

$quantiles.introgression: Quantiles of introgression proportions across individuals.

$sd.introgression: Standard deviation of introgression proportions across individuals.

---

#### create.constant *Create Constant Population Sizes*

---

##### Description

This function allows a carrying capacity (maximum number of individuals a deme/population can sustain) to be specified for genetic simulation using evolve2.0. Specifically, using the argument constant, one can keep sizes of select populations at or below a user-defined threshold.

##### Usage

```
create.constant(pop.index, pop.size)
```

##### Arguments

| | |
|---|---|
| pop.index | *numeric* A number or vector specifying the indices of populations. |
| pop.size | *numeric* A number or vector specifying the size(s) to which each population in pop.index should be kept equal to or below. This argument should be of same length as pop.index and follow the same order - first values of both arguments are used together. |

#### create.events — *Create Historical Events*

##### Description

This function generates the data frame needed to switch between pre-defined migration matrices at select generations while running `evolve2.0`.

##### Usage

```
create.events(nb.events, generation, migration.rate.switch, write = F)
```

##### Arguments

| | |
|---|---|
| `nb.events` | *numeric* The number of historical events (the number of times migration matrices would like to be switched). |
| `generation` | *numeric* Generation(s) you wish to change the migration rate between populations. |
| `migration.rate.switch` | *numeric* The index (number) of the matrix you wish to use at a select generation (see `evolve2.0` for details). |
| `write` | *logical* If true, the events file will be saved to the working directory. |

##### Value

This function returns a data frame with two columns (Generation and migration.rate) and a number of rows equal to the number of historical events. The column 'Generation' records the generation at which one want to modify the migration between populations. The column 'migration.rate' records which of the migration matrices would like to used from that generation forward. See `evolve2.0`.

#### create.source — *Create a Source Population*

##### Description

This function allows you to create a `source` population for genomic simulation using `evolve2.0`. Specifically, using the argument `source`, one can define a population serving indefinitely as a source of individuals. This function generates the value that needs to be fed to the `source` argument in `evolve2.0` if such a demographic scenario wishes to be simulated.

##### Usage

```
create.source(N, nl, pop.index, allele.code)
```

##### Arguments

| | |
|---|---|
| `N` | *numeric* Number of individuals in the source population. |
| `nl` | *numeric* Number of simulated loci. |
| `pop.index` | *numeric* An integer giving the index of the population that wishes to be used as source in the initial `struct` object (obtained using `initial.struc` or `homogeneous.struct`) |
| `allele.code` | *numeric* An integer specifying how the allele (fixed within the source population, see `homogeneous.struct`) should be encoded. |

---

`evolve2.0` *Evolution of Genetic and Genomic Landscape of Introgression:* `evolve2.0`

---

##### Description

This function is identical to `evolve` in `glads`, except that it allows to vary (change) the migration rates over time by defining a data frame describing the desired migration rate at a select generations (see `create.events`). It also allows (1) a population to serve as a recurrent source of individuals (see `create.source`) and (2) populations to have various carrying capacities (maximum number of sustainable individuals; see `create.constant`). Finally, this functions allows the retrieval of simulated data at user-defined generations to evaluate temporal changes in population genetic structure.

##### Usage

```
evolve2.0(
  x,
  time,
  type = c("constant", "dynamic", "additive", "custom"),
  recombination = c("map", "average"),
  recom.rate,
  loci.pos = NULL,
  chromo_mb = NULL,
  init.sex = NULL,
  migration.rate.list = NULL,
  migration.rate.initial = NULL,
  mutation.rate = NULL,
  param.z = NULL,
  param.w = NULL,
  fun = c(phenotype = NULL, fitness = NULL),
  events = NULL,
  gen.snapshot = NULL,
  source = NULL,
  logfile = FALSE,
  constant = NULL
)
```

##### Arguments

`migration.rate.list`
*list* A list of square matrices defining the migration rate between all populations simulated. The index of these matrices (1,2,3,...) is used in the events file (see `create.events`) to change the migration rate at desired generations.

migration.rate.initial
: *numeric* The index of the matrix that wishes to be used at the beginning of simulation (until the occurrence of the first historical event).

events
: The events data frame generated using create.events function.

gen.snapshot
: *numeric* A vector of generations at which you want data to be output (saved).

source
: *list* A list describing the structure of the source population generated using create.source.

logfile
: *logical* Whether or not a log file recording historical events should be generated (TRUE = yes, FALSE = no).

constant
: This option allows to keep the size of select populations at or below a user-defined threshold by randomly discarding the excess of individuals. Takes as input create.constant.

#### See Also

evolve for additional details on the function arguments.

---

extract_popsize *Extract Sizes of Populations from Simulation Results Returned by iterate_evolve2.0.*

---

#### Description

This function allows to extract the size of all populations from an object return by iterate_evolve2.0. Note that this function can only be used when generation snapshots are taken (see help for evolve2.0 for details).

#### Usage

```
extract_popsize(
  data,
  save.details.rds = FALSE,
  out = "extract_popsize_details.rds"
)
```

#### Arguments

data
: A R object or the path to a .rds file containing genomic simulation results.

save.details.rds
: *logical* Whether a list with population sizes recorded at each generation for every population across iterations (n_rep, see iterate_evolve2.0 for details) should be saved in .rds format.

out
: *character* If save.details.rds is TRUE, then rename the output file. Default is "extract_popsize_details.rds".

`homogeneous.struct` *Create Homogeneous Initial Population Structure*

#### Description

This function allows to set an initial population structure for which all individuals within populations are fixed for a given allele (see the help for `initial.struct` for more details).

#### Usage

```
homogeneous.struct(N, nl, n_pop = 1, allele.code = 2)
```

#### Arguments

| | |
|---|---|
| N | *numeric* Number of individuals within (the) initial population(s). |
| nl | *numeric* Number of simulated loci. |
| n_pop | *numeric* Number of populations with the same initial structure to generate. |
| allele.code | *numeric* The code to give to the allele fixed within (the) population(s). |

`iterate_evolve2.0` *Wrap up function: Perform Multiple Simulations*

#### Description

These functions allow the user to run a user-defined number of simulations using `evolve2.0`.

#### Usage

```
iterate_evolve2.0(
  x,
  time,
  type = c("constant", "dynamic", "additive", "custom"),
  recombination = c("map", "average"),
  recom.rate,
  loci.pos = NULL,
  chromo_mb = NULL,
  init.sex = NULL,
  migration.rate.list = NULL,
  migration.rate.initial = NULL,
  mutation.rate = NULL,
  param.z = NULL,
  param.w = NULL,
  fun = c(phenotype = NULL, fitness = NULL),
  events = NULL,
  gen.snapshot = NULL,
  source = NULL,
  logfile = FALSE,
  constant = NULL,
  n_rep = 100,
  nb.cores = 1
)
```

#### Arguments

| | |
|---|---|
| `n_rep` | *numeric* Number of simulations to be performed using `evolve2.0`. |
| `nb.cores` | *numeric* Number of cores to use for simulation. |

#### Note

To extract sizes of populations simulated, use `extract_popsize`.

#### See Also

For details on other arguments see `evolve2.0` `evolve`.

### Index

compute_introgression_lengths, 1
compute_introgression_peaks, 2
compute_introgression_proportions, 3
create.constant, 4, *5*
create.events, 4, *5*
create.source, *5*, 5

evolve2.0, *2*, *4*, *5*, 5, *7*, *8*
extract_popsize, 7, *8*

homogeneous.struct, 7

initial.struct, *7*
iterate_evolve2.0, *7*, 8
