## Appendix S4 for "Temporal variation in introgressed segments’ length statistics sheds light on past admixture pulses": Companions4glads_Example.R

```
#----------------------------#
# Example - companions4glads #
#----------------------------#

# This script provides examples for the implementation of functions compiled
# within the "companions4glads" packages. Below is the code necessary for 
# simulating a two-population system with two bouts of introgression from population 
# 1 into population 2, as well as estimating all three summary statistics:
# (1) introgression proportions (PI), (2) lengths of introgressed sequences (LIS), 
# and (3) within-individual variation in introgressed sequences' lengths (VIS).

#------------------------------------------- Example: Simulations -------------------------------------------#

# Below is the code to simulate the multiple-pulse model described in the main text with a migration (admixture)
# rate of 0.03. 

# To install companions4glads: install.packages("companions4glads_1.2.tar.gz", repos = NULL)
#   !!! Additional packages may need installing !!!

# To install glads: devtools::install_github("https://github.com/eriqande/glads.git")

require(companions4glads)

pop1 <- homogeneous.struct(N = 100, nl = 200, n_pop = 1, allele.code = 1)
pop2 <- homogeneous.struct(N = 100, nl = 200, n_pop = 1, allele.code = 2)

events <- create.events(nb.events = 3,
                        generation = c(6, 201, 206),
                        migration.rate.switch = c(1, 2, 1))

migration.rate.list <- list(matrix(c(0,0,0,0),2,2),
                            matrix(c(0,0.03,0,0),2,2))

chromo_mb <- 150000000
loc.pos <- unlist(lapply(1:200, function(i){i*chromo_mb/200}))

set.seed(5)
multiple.migration <- evolve2.0(x = list(pop1,pop2), 
                                time = 1500,
                                type = "dynamic",
                                param.z = list(list(0.5), list(0.5)),
                                param.w = list(list(4,0.02), list(4,0.02)),
                                recombination = "average",
                                recom.rate = 1.2e-8,
                                loci.pos = loc.pos,
                                chromo_mb = chromo_mb,
                                init.sex = NULL,
                                migration.rate.list = migration.rate.list,
                                migration.rate.initial = 2,
                                mutation.rate = NULL,
                                events = events,
                                gen.snapshot = c(seq(1,20,2), seq(30, 190, 20), seq(192, 300,2), seq(400, 1400, 100)),
                                source = NULL,
                                logfile = FALSE,
                                constant = NULL)
str(multiple.migration)

#------------------------------------------- Example: Estimation --------------------------------------------#

# The code below estimates all three genomic summary statistics evaluated throughout the manuscript for one 
# simulation of the multiple-pulse model with an admixture rate of 0.03. Plots generated at the end mimic 
# Figure 2, 3, and 4 of the manuscript for a single simulation (Figures are based on 100 simulation replicates).

###
# Proportions of introgression
###
prop.int <- lapply(multiple.migration, function(a){compute_introgression_proportions(x = a,
                                                                                     pop = 1,
                                                                                     introgressed.allele = 2,
                                                                                     loc.pos = loc.pos)$mean.introgression})
head(prop.int)
data.introgression <- data.frame(generation = gsub("generation_", "", names(unlist(prop.int))),
                                 introgression = unlist(prop.int)); rownames(data.introgression) = NULL
head(data.introgression)

###
# Length of Introgressed sequences
###
len <- lapply(multiple.migration, function(a){mean(compute_introgression_lengths(x = a,
                                                                                 pop = 1,
                                                                                 allele = c(1,2),
                                                                                 loc.pos = loc.pos,
                                                                                 stats = NULL,
                                                                                 exclude = F,
                                                                                 pool = T)$nb.loci.pool)})
head(len)
data.lengths <- data.frame(generation = gsub("generation_", "", names(unlist(len))),
                           lengths = unlist(len)); rownames(data.lengths) = NULL
head(data.lengths)

###
# Within-individual variation in introgressed sequences's lengths
###
var <- lapply(multiple.migration, function(a){compute_introgression_lengths(x = a,
                                                                            pop = 1,
                                                                            allele = c(1,2),
                                                                            loc.pos = loc.pos,
                                                                            stats = "nb.loci",
                                                                            exclude = F,
                                                                            pool = F)$mean})
head(var)
data.var <- data.frame(generation = gsub("generation_", "", names(unlist(var))),
                       variation = unlist(var)); rownames(data.var) = NULL
head(data.var)

###
# Graphical representation
###
par(mfrow=c(3,1))
plot(introgression ~ generation, data = data.introgression, type = "p", col = "red", pch = 19,
     xlab = "Generation", ylab = "Average proportion of introgression (PI)", ylim = c(0,1))
lines(introgression ~ generation, data = data.introgression, col = "red")
plot(lengths ~ generation, data = data.lengths, type = "p", col = "blue", pch = 19,
     xlab = "Generation", ylab = "Average length of introgressed sequences (LIS)", ylim = c(0,200))
lines(lengths ~ generation, data = data.lengths, col = "blue")
plot(variation ~ generation, data = data.var, type = "p", col = "forestgreen", pch = 19,
     xlab = "Generation", ylab = "Average within-individual variation in introgressed sequences' lengths (VIS)", ylim = c(0,100))
lines(variation ~ generation, data = data.var, col = "forestgreen")
```
