## Appendix S4 for "Temporal variation in introgressed segments’ length statistics sheds light on past admixture pulses": readme.txt

### READ ME ###

This directory contains the compiled R package used in our study to conduct simulations and to estimate introgression summary statistics (companions4glads_1.2.tar.gz).
The manual describing every function within the packages and how to use them is also provided (companions4glads_1.2.pdf). Help for every function may also be accessed
within R using either "?function_name" or "help(function_name)".

Finally, a R script examplifying the use of the package to replicate results presented in the main text is provided (Companions4glads_Example.R).
